# Condensin I confers Stiffness and Centromeric Cohesion to Mitotic Chromosomes

**DOI:** 10.1101/2025.04.29.651176

**Authors:** Christian F. Nielsen, Hannes Witt, Andrea Ridolfi, Bas Kempers, Emma M.J. Chameau, Stijn van der Smagt, Marin Barisic, Erwin J.G. Peterman, Gijs J.L. Wuite, Ian D. Hickson

## Abstract

When cells divide, the newly replicated sister chromatids must be segregated evenly to the daughter cells. During mitosis, mechanical force is applied by spindle microtubules in 2 ways: first by pushing on chromosome arms to promote chromosome congression to the cell equator in metaphase, and then by pulling on kinetochores to promote sister chromatid disjunction during anaphase. For segregation to proceed faithfully, the pliable interphase chromatin must be transformed into stiff mitotic chromosomes able to withstand these forces. However, it is unclear how the cell establishes chromosome stiffness and what the consequences are for dividing cells if this stiffness is disrupted. Many of the structural changes imposed on chromosomes in mitosis are driven by Condensin complexes, in conjunction with Topoisomerase IIα. Here, we have combined rapid protein depletion and live cell imaging with in-depth mechanical characterization of purified mitotic chromosomes to probe the roles of Condensins I and II in the establishment and maintenance of the mechanical strength of mitotic chromosomes. We show that Condensin I, but not Condensin II, is required to establish chromosome stiffness and chromatin elasticity, and yet ceases to be required for the maintenance of these properties once chromosome formation has been completed. Nevertheless, depletion of Condensin I from already formed chromosomes still impacts centromeric chromatin and leads to a loss of sister centromere cohesion. We propose that the extensive chromatin loop network established by Condensin I is locked in place by Topoisomerase IIα mediated DNA catenation.

## Main

During mitotic cell division, the newly-replicated DNA is condensed and resolved to individualized chromosomes that are aligned along the equator of the cell in metaphase. Removal of sister chromatid cohesion, then allows the liberated sister chromatids to be segregated to the two daughter cells in anaphase and telophase. Errors in chromosome segregation can lead to loss or gain of whole chromosomes and the generation of aneuploid cell progeny. Aneuploidy has been implicated as a driver of cancer and several congenital disorders^1^. The protein machinery that aligns and segregates chromosomes is the mitotic spindle, which exerts two types of force on mitotic chromosomes. First, polymerizing spindle microtubules exert polar ejection forces that push on whole chromosomes to promote chromosome congression. Second, each kinetochore is subjected to pulling forces that are applied once the microtubules are attached in a bioriented manner to sister kinetochores. These spindle forces respectively drive chromosomes to the spindle equator in metaphase and pull sister chromatids toward the spindle poles during anaphase/telophase. In addition to these microtubule-mediated forces, mitotic chromosome compaction promoted by histone deacetylation^2^ and polyvalent cations and polyamines^3^ expels water from chromosomes and increases the osmotic force exerted on the chromosomes. Finally, a shearing force is applied when chromosomes are moved through the highly viscous cytoplasm, although, this viscosity decreases when cells enter mitosis^4^. Mitotic chromosomes must, therefore, acquire adequate structural stiffness and elasticity to be able to resist these forces, and hence be aligned and segregated correctly.

The dramatic metamorphosis of chromosomes during mitosis is driven by a combination of histone deacetylation-dependent chromatin compaction^2,3^, and an elaborate sculpting of chromosomes by the concerted action of Condensins I and II in collaboration with Topoisomerase IIα (TOP2A) (reviewed in ^5,6^). The Condensin I and II complexes belong to the Structural Maintenance of Chromosomes (SMC) family that also includes Cohesin and the SMC5/6 complex. Condensins I and II share core SMC2 and SMC4 components, but differ in their non-SMC subunits, with Condensin I including NCAPH, NCAPD2 and NCAPG, and Condensin II NCAPH2, NCAPD3 and NCAPG2^7^. Condensins I and II also differ in their subcellular localization throughout the cell cycle: Condensin II is present on chromatin throughout interphase and mitosis, while Condensin I is largely excluded from the nucleus in interphase until the time of nuclear envelope breakdown in prometaphase^8^. The first stage of mitotic chromosome formation is correspondingly driven by Condensin II and TOP2A in the mitotic prophase^8,9^. In this process, which also depends on prophase pathway mediated removal of Cohesin from chromosome arms^10^, chromosomes are individualized and entanglements between sister chromatid arms are resolved^9^.

The function of both Condensins during mitosis appears to derive from their ability to catalyze extensive DNA loop extrusion^11^. To promote structural changes to chromosomes in mitosis, Condensin complexes are suggested to form a central scaffold from which the chromatin fiber emanates in a series of radial loops^12–14^. To ensure efficient packing of the chromatin fiber, it has been proposed that Condensin complexes form nested loops, with large loops of approximately 400 kb being formed by Condensin II, which are then subdivided into loop domains of around 80 kb by Condensin I^12^. In support of this, chromosomes from Condensin I depleted cells are generally reported to be shorter and wider, following a prolonged prometaphase arrest, consistent with longer loops, while those from Condensin II depleted cells are longer and thinner, consistent with shorter loops^7,15,16^.

Live-cell imaging studies have shown that the combined depletion of Condensins I and II^17,18^ leads to a loss of chromosome structural integrity. This in turn weakens the ability of chromosomes to resist mitotic spindle forces, leading to deformation of centromeric chromatin and misalignment of chromosomes^2,17^. These data are consistent with mechanical assays using micropipettes demonstrating that long-term depletion of both Condensins from human cells reduces the elastic stiffness of mitotic chromosomes^19^. Thus far, however, a comprehensive study of how the individual Condensin I and II proteins shape mitotic human chromosomes to resist mitotic spindle forces is lacking.

In this study, we have investigated the roles of Condensin I and II in both the establishment and maintenance of the structural integrity of human chromosomes and their ability to resist different types of force. For this, we have combined live-cell imaging following rapid depletion of either Condensin I or II using an auxin inducible degron (AID) system with mechanical characterization of native and Condensin-depleted mitotic chromosomes using atomic force microscopy and optical tweezers. We show that Condensin I, but not Condensin II, is required for the establishment of global chromosome stiffness and stability. By varying the timing of Condensin I depletion, we reveal that Condensin I is no longer required for maintenance of the global structural integrity of already condensed chromosomes. Nevertheless, Condensin I is still crucial for the maintenance of centromeric chromatin and cohesion in metaphase and, thereby, for resistance to spindle pulling forces.

## Results and Discussion

### Probing the local properties of the mitotic chromatin network

For mitotic chromosomes to properly resist and respond to the polar ejection and lateral pulling forces of the mitotic spindle (Fig. 1a), they must achieve an appropriate level of elasticity and stiffness. To assess the contribution of Condensins to the establishment of the necessary mechanical properties of mitotic chromosomes, we employed an auxin-inducible degron system (AID) to rapidly deplete either the NCAPH subunit of Condensin I (ED Fig 1a-e) or the NCAPH2 subunit of Condensin II^20^ (ED Fig. 1f). Because mitotic chromosomes continue to undergo hyper-condensation during a prolonged arrest in prometaphase, we sought to achieve tight temporal control of cell cycle kinetics. To this end, we modified Condensin-AID cell lines using a chemical genetics approach^21,22^ to generate cell lines that could be arrested in late G2 phase and released in a synchronous manner (Methods, ED Fig. 1g-j). Cells were depleted of either Condensin I or II during this G2 arrest and then released in the presence of nocodazole and auxin. Native chromosomes were then purified from the resulting prometaphase-arrested cells^23^ (Fig 1b, Methods).

**Figure 1.**
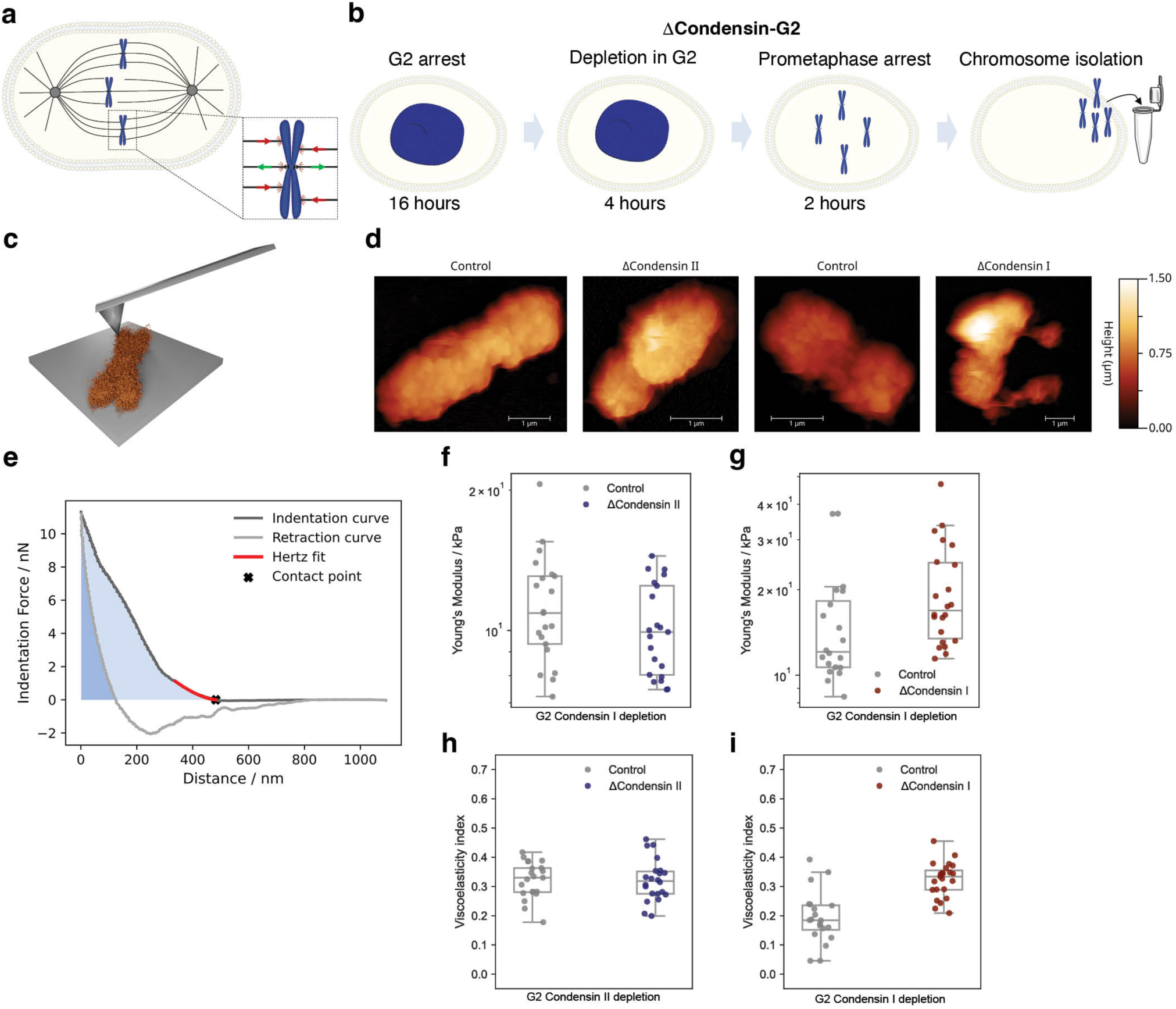
Local structural and mechanical properties of Condensin depleted chromosomes. **a** Simplified diagram of how the mitotic spindle applies polar ejection forces on chromosomes (red arrows) and pulling force on centromeres of chromosomes (green arrows). **b** Scheme for pre-mitotic depletion of Condensins. Cells were arrested in G2 (CDK1as) for 16h and depleted of Condensin I or II in the final 4h (ΔCondensin G2), released and arrested in prometaphase (nocodazole) for 2h before chromosome isolation. **c** Schematic of the AFM indentation of mitotic chromosomes. **d** Representative AFM images of native isolated chromosomes. **e** Representative force distance curve of a chromosome indented by AFM, the red line indicates the Hertz fit, the viscoelastic index is calculated from the area between the indentation curve *A_indent_* (areas shaded in light and dark blue combined) and the area between the retraction curve *A_retract_* (shaded in darker blue) in the loading and unloading regimes, as 𝜂 = 1 − *A_retract_*/*A_indent_* **f,g,h,i** Young Modulus (**f**,**g**) and viscoelastic index (**h**,**i**) of chromosomes depleted of Condensin I (**f**,**h**) or II (**g**,**i**) as derived from AFM experiments.

In agreement with previous studies, the Condensin II depleted chromosomes (henceforth, ΔCII-G2 chromosomes) were slightly longer and thinner compared to control chromosomes (ED Fig. 1k-l, p<0.0001)^7,15,16^. The Condensin I depleted chromosomes (henceforth, ΔCI-G2 chromosomes) appeared less structured with a less defined sister chromatid border, but were otherwise morphologically similar to control chromosomes (ED Fig. 1 m,n). Nevertheless, when cells were instead held for a longer period in prometaphase (5 hours after release from G2 rather than 2 hours) following depletion of Condensin I, chromosomes were shorter and wider, as has been observed in previous studies (ED Fig. 1n-o, p<0.0001). This suggests that the iconic short/wide chromosome morphology observed following Condensin I depletion^7,15,16^ only becomes apparent following a prolonged prometaphase arrest when hyper-condensation of chromosomes takes effect. To avoid this potentially confounding issue, we released cells from G2 arrest for only 2 hours in subsequent chromosome analyses.

To inspect the chromosome surface topology in greater detail, we imaged native and Condensin-depleted chromosomes using atomic force microscopy (AFM, Fig. 1c). The surface of control and ΔCII-G2 chromosomes were characterized by an almost identical “granular” textured structure reminiscent of the texture of mitotic chromosomes found previously^24^. By contrast, the ΔCI-G2 chromosomes had a less structured surface with protrusions and decreased granularity. The less structured surface made it harder for the AFM tip to accurately follow the chromosome topology, decreasing resolution specifically for ΔCI-G2 chromosomes (Fig. 1d). We speculated that this could be caused by fewer crosslinks and loop arrays in the chromatin structure,

To further probe this and how Condensins might contribute to the resistance of chromosomes to the pushing force of the spindle microtubules during chromosome congression, we utilized AFM-based Force Spectroscopy (AFM-FS). AFM-FS can impart localized nano-indentations into chromosomes to mimic the pushing force used by microtubules to promote chromosome congression^25,26^. The resulting force curves (Fig. 1e) describe the relationship between the indenting force and the tip-sample distance, and permit the characterization of the chromatin network’s local mechanical response^26^. By fitting the initial part of the force curves with a modified Hertz model (Fig. 1e), we extracted the Young’s Modulus^27^, which describes the stiffness of the probed chromatin network (Fig. 1f,g). Interestingly, neither the ΔCII-G2 (Error is SEM, Control: 11.4 ± 0.7 kPa, ΔCII-G2: 10.2 ± 0.5 kPa, p=0.16) (Fig. 1f) nor ΔCI-G2 (Error is SEM, Control: 16 ± 2 kPa, ΔCI-G2: 20 ± 2 kPa, p=0.09) (Fig. 1g) chromosomes showed a significant difference in Young’s Modulus compared to control chromosomes (ED Table 1). By measuring the hysteresis between the indentation and the retraction curves (see Methods), we calculated the viscoelasticity index (η), which quantifies the relative contribution of viscosity and elasticity to the overall mechanical response; in essence, the ability of the chromosomes to elastically recover from applied indentations^28^ (Fig. 1h,i). Values of η close to 0 correspond to a fully elastic response, while values close to 1 indicate that all the indentation energy is viscously dissipated by the sample. There was no discernible difference between the responses of ΔCII-G2 and control chromosomes (Error is SEM, Control: 0.32 ± 0.01, ΔCII-G2: 0.32 ± 0.02, p=0.98) (Fig. 1h, ED Table 1). By contrast, ΔCI-G2 chromosomes had a significantly higher viscoelasticity index than the respective controls, indicative of a more viscous response of the chromatin network (Error is SEM, Control: 0.19 ± 0.02, ΔCI- G2: 0.32 ± 0.01, p=1.3*10^-6^) (Fig. 1i, ED Table 1). The observed difference would be consistent with a reduction in cross-linking of the chromatin network in cells lacking Condensin I, which would generate chromosomes of a more viscous nature and less prone to recover from deformations.

It was established recently that Condensin proteins are not essential for mitotic chromosomes to resist piercing by spindle microtubules^2^. Our finding that Condensin depletion does not affect the local stiffness of the chromosome surface is consistent with that finding. Resistance to piercing in cells is, instead, governed by a charge-dependent repulsion of microtubules in the surface compartment of chromosomes at least in part due to deacetylation of chromatin^2^. Nevertheless, previous studies have indicated that Condensins are still required for proper congression of chromosomes in metaphase^8,29^. During congression, the chromatin network must be sufficiently elastic to resist local deformation from the force applied by the microtubules. We suggest that the congression defect of ΔCI-G2 chromosomes might, therefore, be explained by the increased viscosity of the chromatin network as observed here for mitotic chromosomes established without Condensin I.

### The global mechanical properties of mitotic chromosomes

Next, we analyzed the contribution of Condensins to the establishment of global mitotic chromosome stiffness. For this, we attached streptavidin-coated microspheres to chromosomes biotinylated on their telomeres (Fig. 2a,b) and then applied force along their longitudinal axis using optical tweezers (OT)^23^. Consistent with our previous studies^3,23^, we observed that human chromosomes display a characteristic non-linear stiffening. This general feature of the chromosomes was not affected by depletion of either Condensin enzyme (ED Fig. 2a and Fig. 2c,d). Nevertheless, analysis of mean stretching curves revealed that there was a higher degree of deformation of ΔCI-G2 chromosomes at any given force compared to control chromosomes (Fig. 2c). In contrast, while ΔCII-G2 chromosomes were generally longer than control chromosomes, their deformation as a function of force was indistinguishable from that of the controls (Fig. 2d). The distinct response of ΔCI-G2 chromosomes became more apparent when plotting the stiffness of the chromosomes as a function of force applied (Fig. 2e-f). For each individual stretching curve, we extracted the linear stiffness (ED Fig. 2f), the power law exponent (ED Fig. 2g) and the critical force (ED Fig. 2h), which provide a more detailed mechanical description of the chromosome response^23^. This analysis confirmed that there was a strong reduction in the linear stiffness at low forces below approximately 100 pN after Condensin I depletion (Error is SEM, Control: 71±14 pN/µm, ΔCI-G2: 21 ± 3, *p* < 10^-4^,Fig. 2e, ED Fig. 2d), but not after Condensin II depletion (Error is SEM, Control: 64±17 pN/µm, ΔCII-G2: 49 ± 15, *p* = 0.6, Fig. 2f, ED Fig. 2e, ED Table 1). These data indicate that depletion of Condensin I before mitosis generates more pliable chromosomes that are less able to resist force. We conclude that Condensin I is crucial for establishing the appropriate stiffness of mitotic chromosomes, whereas Condensin II is not. To assess how the depletion of Condensin I or II impacts the global viscoelastic properties of chromosomes, we performed oscillatory optical tweezer experiments^23^. In this case, depletion of neither Condensin I nor Condensin II had a measurable impact on the global fluidity of chromosomes over a wide range of frequencies (ED Fig. 2g-j). This shows that, although the fluidity of the local chromatin network is increased in ΔCI chromosomes, when chromosomes are probed globally by longitudinal force oscillations, global fluidity is not affected by loss of either Condensin.

**Figure 2.**
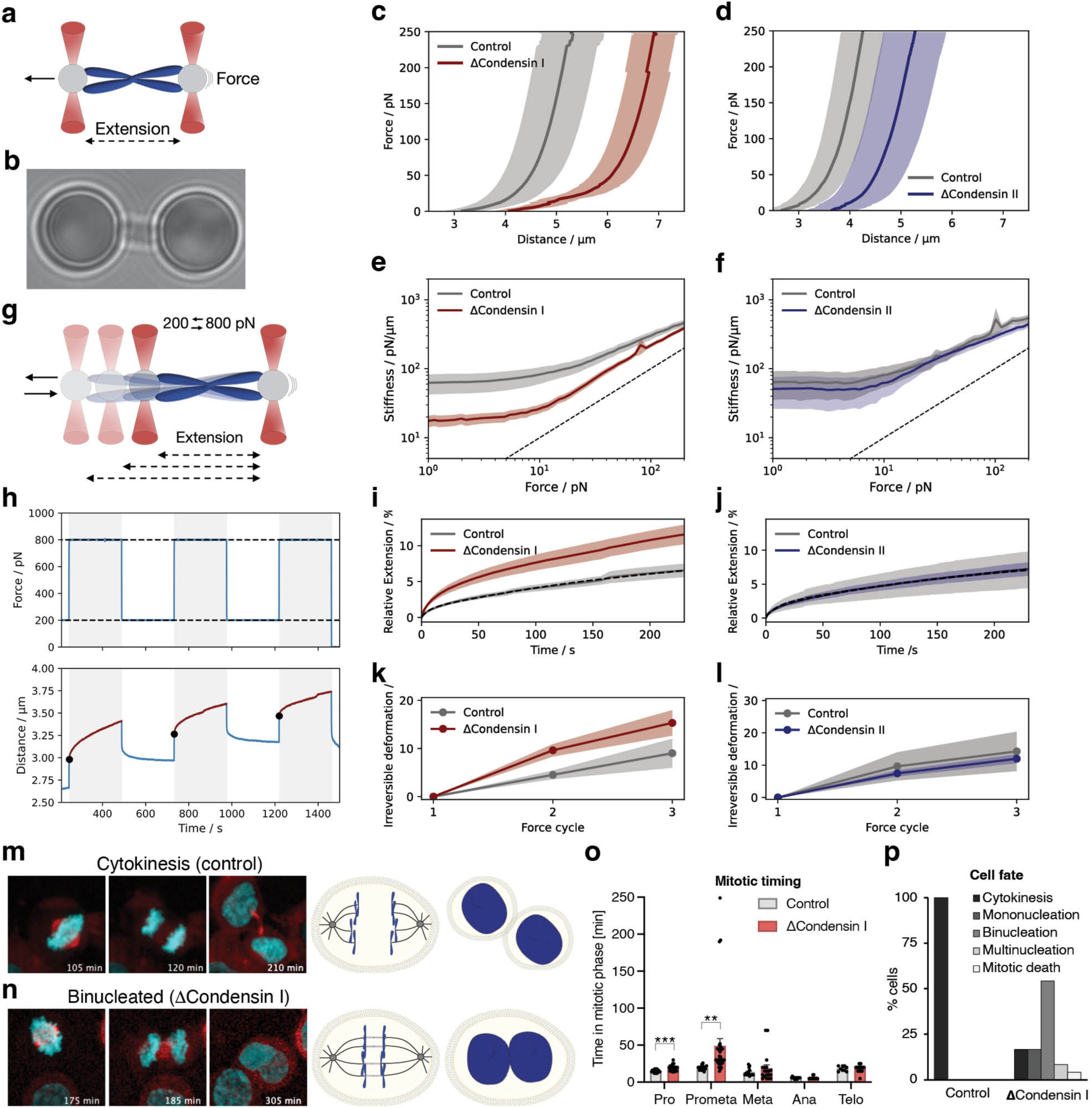
Global structural properties of ΔCondensin chromosomes **a** Diagram and **b** image of a chromosome clamped between two microspheres held by optical tweezers. **c**,**d** Average stretch curves for ΔCondensin I (**c**) and II (**d**) chromosomes. **e,f** Stiffness as a function of force for ΔCondensin I (**e**) and II (**f**) chromosomes, as derived from optical tweezers experiments. **g** Diagram and **h** scheme of optical tweezers-based stability assay. The force is alternated between 800 and 200 pN in consecutive cycles. The relative extension at high force (shading in **h**) as well as the length change between the beginning of consecutive high force cycles (black circles in **h**) to assess plastic deformation was analyzed. **i,j,k,l** Mean relative extension at high force (**i,j**) and mean relative plastic deformation (**k,l**) of ΔCondensin I (**i**,**k**) and II (**j**,**l**) chromosomes. Shaded areas in **c, d, e**, **f**, **i, j**, **k** and **l** are 1.386 times the SEM, such that touching error bars represent a significance level of 95%. **m,n,o,p** Cells treated like in Fig. 1b were imaged following release from G2 arrest. Cells express endogenously tagged H2B-EGFP (cyan) to visualize chromatin and ectopically expressed α-tubulin-mCherry (red) for the mitotic spindle. **m** Control cell going through mitosis and cytokinesis. Diagrams depict anaphase and regular G1. **n** ΔCondensin I cell failing mitotic segregation becoming binucleated. Diagrams depict anaphase with unresolved DNA ultrafine DNA bridges (not visible in the anaphase microscopy image) and binucleation. **o** Quantification of the time spent in different mitotic phases. Error bars are SEM. Control n=17, ΔCondensin I n=32. **p** Quantification of cell fate after mitosis. Control n=23, ΔCondensin I n=24. ** p < 0.01 and *** p < 0.001.

So far, we have demonstrated that Condensin I is important for chromosomal stiffness measured by longitudinal stretching of chromosomes at forces up to ∼250 pN. The mitotic spindle, however, has been proposed to exert forces of up to 700 pN^30^. This led us to analyze if Condensins are also important for the mechanical stability of chromosomes subjected to very high force. Mechanical stability can be tested by determining whether the chromosomes deform in a plastic manner during repeated elongation cycles at high force. For this, each chromosome was clamped alternately at very high force (800 pN) and then at a more moderate force (200 pN) for a total of 3 high/moderate force cycles (Fig. 2g,h). We then quantified the degree to which the chromosome was elongated during each cycle of high force stretching. This permitted us to distinguish between viscous creep, which is a reversible process, and permanent deformation of the chromosome. During these high force analyses, the chromosome continuously elongated, without ever reaching saturation, even when held for >30 mins (ED Fig. 3a). We observed that only the ΔCI-G2 chromosomes showed increased elongation compared to controls (Fig. 2i,j and ED Fig. 3b,c and ED Table 2). To quantify permanent plastic deformation, we plotted the difference in length of the chromosomes at the beginning of each high force cycle. This analysis showed that there was a stronger permanent deformation of ΔCI-G2 chromosomes (Fig. 2k) than of either ΔCII-G2 or control chromosomes (Fig. 2l). Hence, Condensin I also provides mechanical stability and resistance to plastic deformation when chromosomes are exposed to high force.

**Figure 3.**
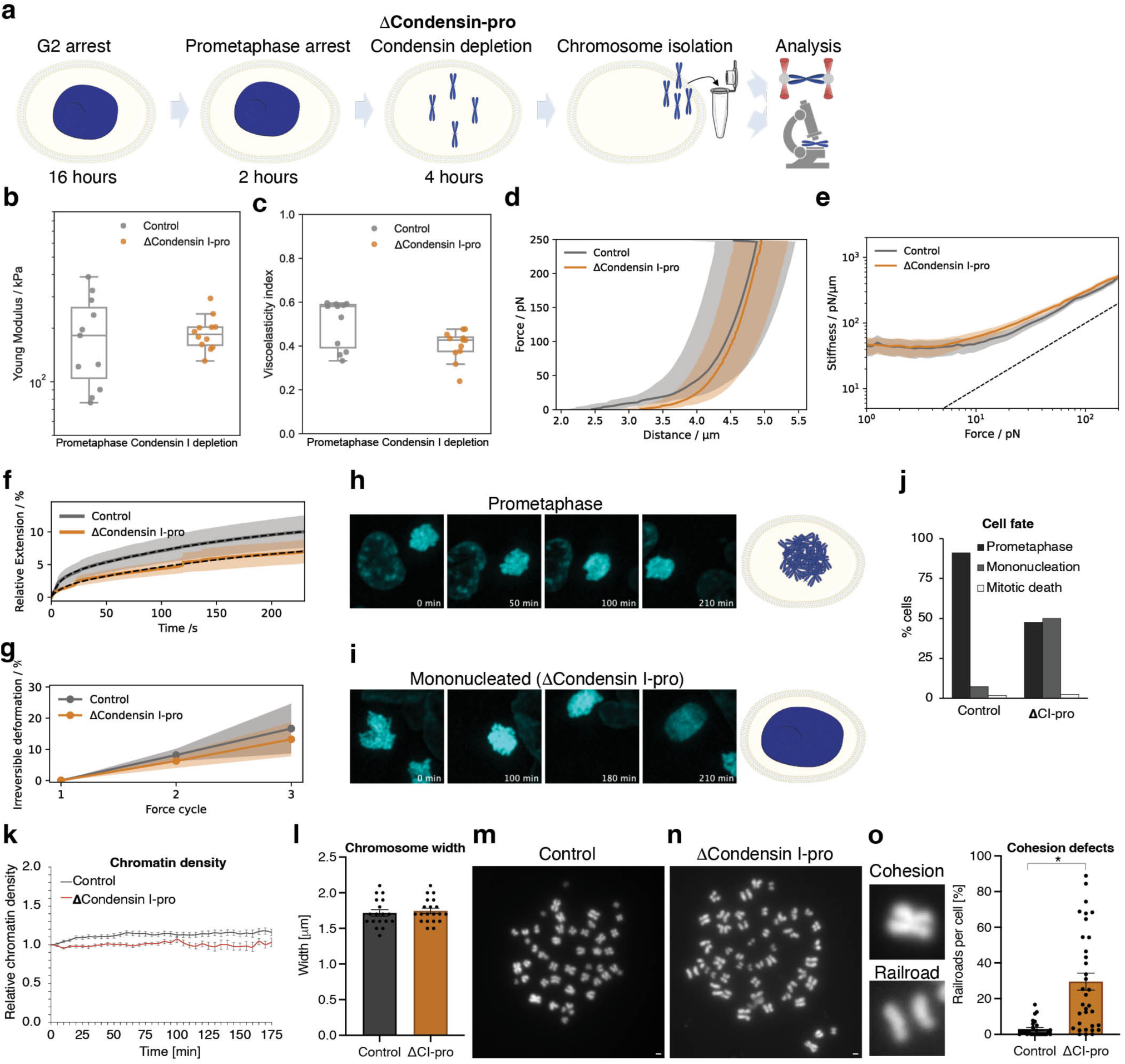
Role of Condensin I in maintenance of mitotic chromosomes **a** Experimental procedure for Condensin I depletion in prometaphase (ΔCondensin I-pro). **b** Young’s Modulus and **c** viscoelastic index of ΔCondensin I-pro chromosomes as derived from AFM-FS experiments. **d** Averaged force distance curves and **e** stiffness as a function of force for ΔCondensin I-pro chromosomes, as derived from optical tweezer experiments. **f** Mean relative extension at high force and **g** mean relative plastic deformation of ΔCondensin I-pro chromosomes. **d**,**e**,**f**,**g** Shaded areas are 1.386 times the SEM such that touching error bars represent a significance level of 95%. **h,i,j** NCAPH-mAID cells treated like in **a** and imaged following 2 h release into nocodazole with or without Condensin I depletion. **h** A control cell in prometaphase and **i** a ΔCondensin I cell in prometaphase becoming mononucleated. **j** Quantification of mitotic cell fate. **k** Quantification of relative chromatin density over time. **l,m,n,o** Representative images (**m,n**) and quantification of chromosome width (**l**) and railroad chromosome cohesion defects (**o**) in chromosome spreads from cells treated like in **a. l**,**m** Each datapoint represents the average of a chromosome spread from one cell. Similar results were obtained from two independent experiments. Error bars are SEM. * p < 0.05.

Next, we considered how the altered mechanical properties of ΔCI-G2 chromosomes might affect chromosome alignment and segregation in mitosis. Therefore, we monitored fluorescent chromatin (histone H2B-EGFP) and microtubules (α-tubulin-mCherry) in control or ΔCI-G2 cells traversing mitosis. While control cells progressed through mitosis with no apparent abnormalities (Fig 2m, ED Fig. 3d, Supplementary Movie 1) ΔCI-G2 cells were slghtly delayed in prophase (19 min compared to 15 on controls, p=0.000024) and very delayed in prometaphase (49 min compared to 20 min in controls, p=0.004) with non-congressed chromosomes (Fig. 2n,o, ED Fig. 3e-f, Supplementary Movie 2-3). Hence, we confirmed previous data showing that chromosome congression is affected by the absence of Condensin I^8,29^. Following exit from metaphase, all control cells analyzed completed cytokinesis (Fig. 2p). By contrast, approximately 75% of the ΔCI-G2 cells underwent a highly abnormal anaphase-telophase where the DNA masses initially moved apart and then appeared to snap together again (ED Fig. 3e,f,g, Supplementary movie 2-4) with more than 80% abandoning cytokinesis (Fig. 2n,p and ED Fig. 3e-h). More than 50% of ΔCI-G2 cells became binucleated in G1 (Fig. 2n,p, ED Fig. 3e, Supplementary movie 2). These results demonstrate that sister chromatids cannot be disjoined properly in anaphase without Condensin I and that the structural properties instilled by Condensin I in mitotic chromosomes are crucial for multiple aspects of mitosis.

### Condensin I is not required for the maintenance of global chromosome stiffness

Condensins have been shown to be very sensitive to even relatively low forces, with measurements showing that loop extrusion ceases at forces above 1 pN^31,32^. Based on this, we deemed it unlikely that Condensin I alone would be capable of ‘crosslinking’ chromatin and conferring stiffness and resistance to mitotic spindle forces. We therefore investigated if Condensin I was still required for the maintenance of mitotic chromosome stiffness after chromosome condensation had been completed. For this, we depleted Condensin I specifically from cells arrested in prometaphase (ΔCI-Pro; Fig. 3a) and analyzed the local and global mechanical properties of these chromosomes. ΔCI-Pro chromosomes did not show a difference in local stiffness (Error is SEM., Control: 192 ± 30 kPa, ΔCI-pro: 190 ± 13 kPa, p= 0.97) (Fig. 3b, ED Table 1) or viscoelasticity index (Error is SEM, Control: 0.50 ± 0.03, ΔCI- Pro: 0.40 ± 0.02, p = 0.07) (Fig. 3c, ED Table 1), as measured by AFM-FS. Similarly, the ΔCI- Pro chromosomes did not show differences in force-dependent lengthening (Fig. 3d) and global linear stiffness (Error is SEM, Control: 44 ± 9, ΔCI-Pro: 51 ± 9, *p* = 0.58, Fig. 3e) or in global fluidity when analyzed using optical tweezers (ED Fig. 4a,b). Surprisingly, these ΔCI- pro chromosomes were not even sensitized to high force elongation (Fig. 3f) or plastic deformation (Fig. 3g). This is in sharp contrast to the findings for pre-mitotic depletion of Condensin I (Fig. 1 and Fig. 2) and indicate that while Condensin I is crucial for the establishment of global mitotic chromosome structure, it appears to be dispensable for the maintenance of that structure once mitotic chromosome formation has been completed.

**Fig. 4.**
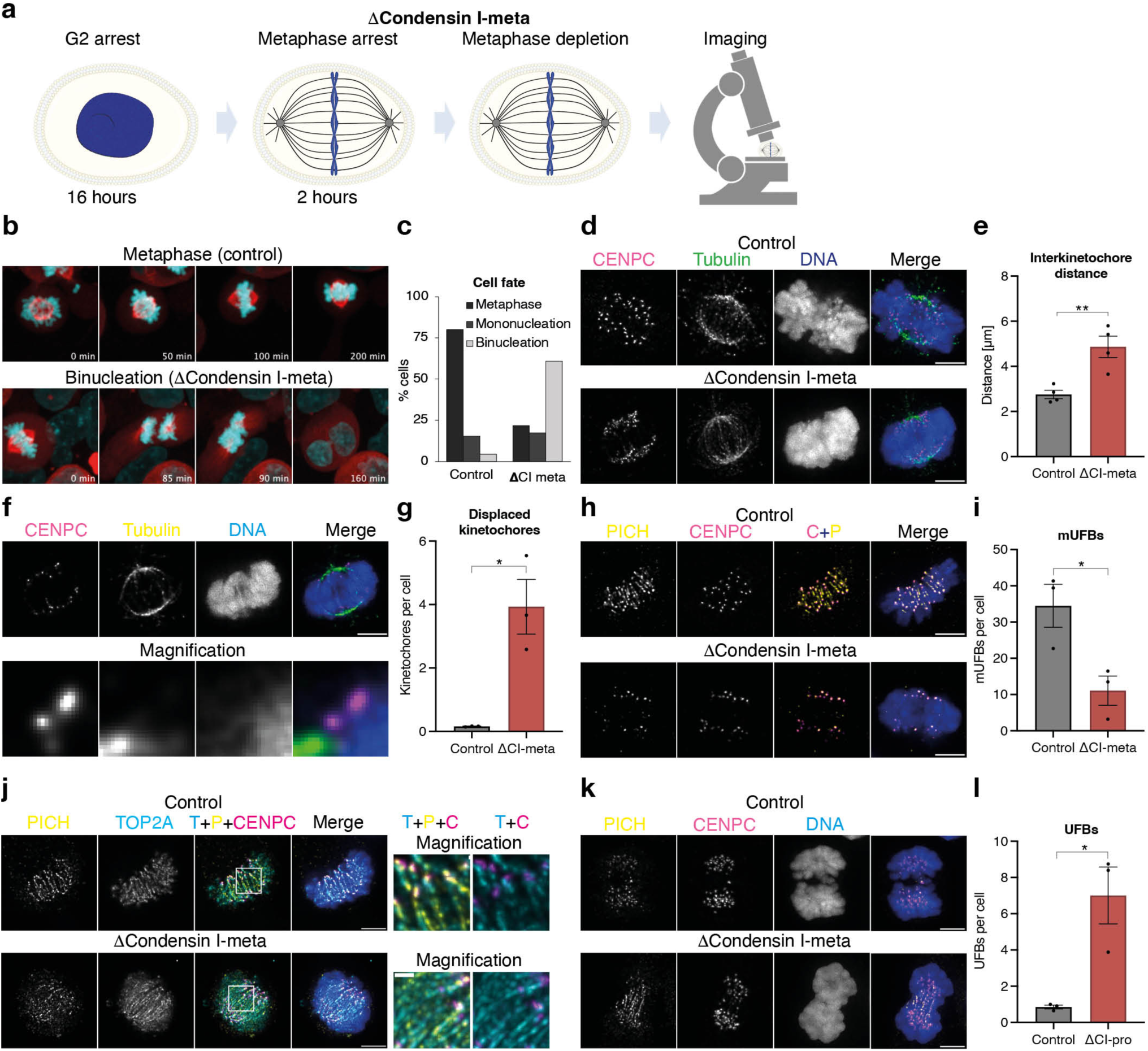
Loss of Condensin I in metaphase with active spindles **a** Diagram of metaphase arrest protocol. NCAPH-mAID cells were arrested in G2 and released into proTAME-Apcin metaphase arrest for 2h, then imaged +/- Condensin I depletion (ΔCondensin I meta). **b** Representative live cell images of a control cell arrested in metaphase (Metaphase) and a cell in metaphase being depleted of Condensin I (Binucleation). Cells express endogenously tagged H2B-EGFP (cyan) to visualize chromatin and ectopically expressed α-tubulin-mCherry (red) for the mitotic spindle. **c** Quantification of cell fate of cells treated and imaged like in **b**. **d,f,h,j** Representative images of cells in metaphase immunofluorescently stained for CENPC (magenta), α-tubulin (green), PICH (yellow) or TOP2A (cyan). DNA is stained with DAPI (blue). The white boxes in **f** and **j** denotes the magnification area. **e**,**g**,**i** Quantification of interkinetochore distance (between CENPC foci, **e**), kinetochores displaced outside chromatin (**g**) and the presence of PICH positive mUFBs (**i**) in metaphase of control or ΔCondensin I-meta cells (2h depletion). **k,l** representative images (**k**) and quantification (**l**) of UFBs in anaphase (**l**) in NCAPH-mAID cells arrested in prometaphase with Nocodazole for 2h, arrested for 2 more hours with and without Condensin I depletion and then released from prometaphase arrest for 45 min with or without Condensin I depletion. DNA (blue) is stained with DAPI. Error bars represent SEM, * p < 0.05 and ** p < 0.01. White scale bars are 5 μm, except in the magnification in **j** where it is 1μm. Datapoints in plots represent the average from independent experiments.

To investigate if Condensin I is required for mitotic chromatin maintenance in cells, we monitored ΔCI-pro cells by live cell imaging. As expected, control cells remained in the prometaphase arrest and showed little change over time (Fig. 3h, Supplementary movie 5). In contrast, ∼50% of the ΔCI-pro cells exited the arrest (∼175 min after addition of auxin; ED Fig. 4c) without attempting anaphase, and underwent cytokinesis to generate mononucleated, but tetraploid, daughter cells (Fig. 3i,j). These data suggest that Condensin I is continuously required to maintain cell arrest after activation of the spindle assembly checkpoint (SAC) by nocodazole treatment. Chromatin density in these prometaphase-arrested cells increased slightly over time in control cells, but did not change following Condensin I depletion (Fig 3k). In line with this, we observed on metaphase spreads that ΔCI- pro and control chromosomes had a similar average width (Fig 3l,m,n). Nevertheless, the ΔCI-Pro chromosomes displayed a clear centromeric cohesion defect (Fig. 3n,o). Indeed, more than 25% of ΔCI-pro chromosomes had lost the characteristic constriction point at the centromere and instead displayed a ‘railroad’ morphology reminiscent of Cohesin-depleted cells, with a 9-fold increase in this aberrant morphology compared to that of control chromosomes (Fig. 3o, p<0.0001). This phenotype was also observed in chromosomes from cells where Condensin I was depleted before mitosis (∼10-fold increase compared to control chromosomes; ED Fig. 4d, p=0.028, and ED Fig. 4e, p=0.0055). These data suggest that Condensin I is continuously required for the maintenance of centromeric cohesion.

### Role of Condensin I in the maintenance of centromeric chromatin

Because of the railroad phenotype discussed above, we hypothesized that a change in centromere structure might be the cause for cells to exit a nocodazole arrest, even though nocodazole drives microtubule depolymerization and loss of spindle force. Hence, we addressed whether the effect of Condensin I depletion might be exacerbated when active spindle forces are maintained during the arrest. This was achieved by arresting cells in metaphase using a combination of proTAME and Apcin^33^, before depleting Condensin I (ΔCI- meta, Fig. 4a). Following such a 5-hour arrest period, 80% of control cells stayed in metaphase with congressed chromosomes (Fig. 4b,c, Supplementary movie 6,7). In contrast, ∼75% of the ΔCI-meta cells entered a premature anaphase and then abandoned cytokinesis (with an average mitotic exit time of ∼100 min after addition of auxin; ED Fig. 4f). Most of the cells undergoing this premature anaphase initially displayed separated chromatin masses before the chromatin was seen to recoil and the cells then abandoned mitosis (Fig. 4b and ED 4g, Supplementary movie 7). The majority of these cells became binucleated (and tetraploid) in the next G1 phase (Fig. 4b,c, Supplementary movie 7). These live cell analyses show that the phenotype of cells lacking Condensin I is significantly exacerbated by the maintenance of an active mitotic spindle and that the SAC cannot be sustained without Condensin I.

We reasoned that, although Condensin I is not required for resistance to longitudinal stretching forces, it might still be necessary for resistance to lateral spindle forces applied to kinetochores, consistent with the enrichment of Condensin I around centromeric chromatin^15,34^ (ED Fig. 4h). Hence, we tested the hypothesis that Condensin I might be necessary for centromeric stiffness in metaphase to resist the force of the mitotic spindle. Consistent with a defect in centromeric stiffness, depletion of Condensin I during metaphase (Fig. 4a) led to a ∼2 μm increase in the inter-kinetochore distance (IKD; the distance between sister CENPC foci; Fig. 4d-e, p=0.0062). In these experiments, Condensin I was depleted for only 2 h to ensure that chromosomes could be analyzed before the cells began to exit mitosis (Fig. ED 4f). It should be noted that the average IKD obtained for our control cells corresponded well with previous studies of cells arrested with active spindles^35^. It was reported previously that depletion of Condensin I^17^ or both Condensins^2,36^ before entry into mitosis leads to a significant increase in IKD. From our ΔCI-meta data, we can conclude that Condensin I continues to be required during mid-mitosis to prevent an anomalous increase in IKD. Moreover, we observed a 24-fold increase in the number of kinetochores (marked by CENPC staining) per cell that were displaced from their usual location within the main chromatin mass. Instead, these displaced kinetochores localized with the mitotic spindle. (Fig. 4f,g, p=0.012). This phenotype was reported previously for cells depleted of both Condensins prior to entry into mitosis^2^. Hence, we conclude that this previously identified role for Condensins at centromeres is conferred by Condensin I.

We confirmed previous data showing that centromeres in metaphase with active spindle forces are frequently connected by short, histone-free, ultrafine DNA bridges (metaphase UFBs; mUFBs), which can be visualized by staining for the DNA translocase, PICH^37^ (Fig. 4h). PICH is recruited to DNA under tension^38^ and is a canonical marker of UFBs^37,39^. Curiously, however, the number of PICH positive mUFBs was significantly decreased in ΔCI- meta cells following only two hours of CI depletion (Fig. 4b-f, mUFBs per cell ± SEM, Control: 35 ± 6, ΔCI-meta: 12 ± 4, p= 0.03). We considered whether this might be explained either by the premature resolution of mUFBs or by a decrease in mUFB tension, leading to a dissociation of PICH. To distinguish between these possibilities, we analyzed the localization of CENPB, which binds specifically to centromeric DNA^40^. We observed that, in the ΔCI-meta cells (and in control cells), the centromeres were connected by thin threads of CENPB- positive centromeric DNA (ED Fig. 4f), suggesting that mUFBs are still present in the absence of Condensin I, but that these mUFBs lack an association with PICH. Strikingly, we observed that the main UFB resolution enzyme, TOP2A, localized strongly to mUFBs in both control cells and ΔCI-meta cells, despite these latter mUFBs lacking PICH. This contrasts with the fact that TOP2A is rarely present on UFBs in anaphase^41^. Taken together, we propose that, in ΔCI-meta cells, tension on mUFBs declines to a point where the affinity of PICH for the UFB is lost.

To investigate whether Condensin I depletion might affect the resolution of UFBs, we analyzed the frequency of PICH-positive UFBs in cells released into anaphase. At this point, tension might be restored to UFBs connecting sister centromeres as the spindle pulls sister chromatids apart, promoting the binding of PICH. Because a proTAME-Apcin mediated arrest in metaphase is irreversible^42^ and because cells depleted of Condensin I before mitotic entry undergo an extended period of arrest in prometaphase (Fig. 2m) and an aberrant anaphase (Fig. 2j), we could not score anaphase UFBs using established protocols. Instead, we adopted a protocol where asynchronously growing cells were treated with nocodazole for 4 hours to arrest them in prometaphase, while depleting Condensin I during the final two hours. Using a modified UFB analysis protocol (see Methods) cells were released from a nocodazole arrest for 45 mins and UFBs were scored in anaphase B cells by immunofluorescent staining for PICH. This revealed a 7-fold increase in the frequency of anaphase UFBs following Condensin I depletion (Fig. 4k,l, p=0.017). These UFBs overwhelmingly originated from centromeres (>95%) (ED Fig. 4g-h). Previous studies reported that siRNA depletion of both Condensins for 72 hours drastically increased the level of UFBs^43^ and that knockout of the Condensin II subunit NCAPH2 increased UFB levels by 3-fold^44^. By comparison, our 2-hour depletion of Condensin I from prometaphase cells increased the frequency of UFBs per cell by 7-fold. Hence, we conclude that Condensin I is required to suppress UFB accumulation in mitosis and that these UFBs are almost exclusively centromeric. The persistence of UFBs in anaphase would also explain why Condensin I-depleted cells become binucleated (Fig.4b,c), because UFBs can prevent the final stage of cell division (abscission), which generates binucleated cells^45^.

Using IKD data to model the centromere as a spring according to Hooke’s law, it was shown previously that Condensins are a determinant of centromere stiffness^36^. The increase in IKD we observe in ΔCI-meta cells, combined with the disappearance of PICH from mUFBs, support this model. Overall, we therefore conclude that although Condensin I is not required for the maintenance of global chromosome stiffness, it is continuously required to support centromeric stiffness.

### General Discussion and Model

We have shown that mitotic chromosomes depend on Condensin I, and not on Condensin II, to achieve chromatin stiffness and to resist structural deformation. Despite this, Condensin I was not continuously required to maintain these properties, because depletion of Condensin I from already compacted mitotic chromosomes did not alter global chromosome stiffness. Instead, we have shown that chromosomes isolated from cells in which Condensin I is removed during metaphase have a sister chromatid cohesion defect likely caused by decreased centromere tension. This leads to dissociation of the DNA tension sensor PICH from UFBs connecting sister centromeres in metaphase. The decreased centromere stiffness makes centromeric chromatin less resistant to the spindle pulling forces, ultimately preventing the resolution of these inter-sister DNA linkages and chromosome segregation. Clearly, our method to probe the mechanical properties of chromosomes by longitudinal stretching is unable to probe the specific properties of the centromere with a lateral organization of its chromatin loops, which would require the development of a method to directly probe the centromere by lateral stretching.

### Why might Condensin II appear to play such a minor role in mitotic chromosomes?

It was unexpected that depletion of Condensin II had no discernible impact on the mechanical properties of chromosomes. In a pioneering micropipette-based study, Condensin II was reported to be an important determinant of mitotic chromosome stiffness^19^. In those experiments, however, Condensin II was depleted with siRNAs over several cell cycles (72 hours), which might have confounding effects, particularly considering that Condensin II is important for genome architecture, transcription and replication in interphase cells^20,46,47^. In our experiments, Condensin II was depleted for only 4 hours from cells already arrested in late G2 phase. Nevertheless, it was still surprising how little impact this had, given the very similar activity of Condensins I and II (loop extrusion) and the fact that both Condensins are localized along the chromosome axis. However, Condensin I is 4 times more abundant than Condensin II on human metaphase chromosomes^13^, most likely increasing the contribution of Condensin I to the formation of the loop network. Based on the nested loop model^12^, Condensin II loop anchors are located at the loop bases and close to the more abundant Condensin I anchors. Therefore, it is conceivable that Condensin II plays little role in providing longitudinal chromosomes stiffness. Consistent with this, hyper-activation of Condensin I has been shown to compensate for Condensin II loss.

### A model for the crosslinking of metaphase chromatin

A recent model posited that a mitotic chromosome acts as phase-separated, gel-like network that is crosslinked by Condensin complexes^6^. In a polymer network such as a mitotic chromosome, crosslinking of the network would confer increased stiffness and solidity. We propose that Condensin I is a more potent contributor to this crosslinking than Condensin II due to its higher contribution to loop generation. Despite this, it was shown previously using single molecule techniques that loop extrusion by Condensin complexes is arrested at forces above 0.5 pN^48^, and that extruded DNA loops dissipate at forces above 10 pN, even though Condensin remains associated with the DNA^49^. This implies that Condensins might be poorly suited by themselves to serve as direct DNA ‘crosslinkers’ that lock loop domains together. These considerations, combined with our finding that loss of Condensin I from already formed mitotic chromosomes does not influence longitudinal chromosomes stiffness, lead us to propose that an alternative factor likely recognizes the chromatin structure established by Condensin I and then locks this structure in place. In that regard, Condensins create DNA structures that are optimal for the action of TOP2A^50–52^. This is an enzyme with an ability to encircle two independent dsDNA segments and to both catenate (entangle) and decatenate (disentangle) dsDNAs through its DNA cleavage and strand passage activity. TOP2A would, therefore, be well suited to serve as a crosslinker to fix the chromatin after loop generation by Condensin I in (pro)metaphase. Consistent with an important role for TOP2A in the maintenance of chromosome integrity, depletion of TOP2A from already condensed metaphase chromosomes leads to chromatin decompaction^53^.

In summary, using a complementary range of biophysical and cellular methods, we have revealed a key role for Condensin I in both the establishment of overall chromosome stiffness and the maintenance of centromere integrity in mitotic human cells. In contrast, we observed no obvious role for Condensin II in conferring stiffness to chromosomes. Our data imply that, even in the absence of the large chromatin loops normally created by Condensin II, there is sufficient time following recruitment of Condensin I to the chromatin for the necessary DNA loop arrangement to be generated that confers stiffness to mitotic chromosomes.

## Acknowledgements

This work was supported by European Union Horizon 2020 grants (Chromavision 665233 to G.J.L.W., I.D.H., and E.J.G.P.; and Antihelix 859853 to I.D.H., and G.J.L.W.), the European Research Council under the European Union’s Horizon 2020 research and innovation program (MONOCHROME, grant agreement no. 883240 to G.J.L.W.), the Novo Nordisk Foundation (NNF18OC0034948 to I.D.H. and G.J.L.W.), the Deutsche Forschungsgemeinschaft (WI 5434/1-1 to H.W.), and the Danish National Research Foundation (DNRF115 to I.D.H.).

## Author Contributions

C.F.N, H.W, E.J.G.P., G.J.L.W., and I.D.H. conceptualized the research, C.F.N. designed and produced chromosomes, performed biochemical characterization. C.F.N. and M.B. performed live cell imaging, H.W., B.K., E.M.J.C., and S.v.d.S. performed optical tweezers experiments, H.W. analyzed optical tweezers data, A.R. performed Atomic Force Microscopy experiments and analyzed the data, C.F.N., H.W., A.R. and I.D.H. wrote the manuscript, and all authors edited it.

## Methods

### Cell culture and cell lines

Cell lines were grown in DMEM with 10% fetal bovine serum (FBS) supplemented with penicillin-streptomycin at 37°C in humidified incubator with 5% CO_2_. The NCAPH-mAID and NCAPH2-mAID cell lines were engineered from the HCT116 TET-OsTIR1 cell line, which was a kind gift from Prof. M. Kanemaki (RIKEN, Japan)^54^. Cloning of the NCAPH2-mAID cell line is described in^20^. For the NCAPH-mAID cell line, the mAID sequence was introduced at the C terminal end of the NCAPH protein by CRISPR-Cas9 directed targeting of the 3’-UTR of the gene using the Zhang lab CRISPR design tool (https://zlab.bio/guide-design-resources; MIT, discontinued). The gDNA sequence was added to the pSpCas9(BB)-2A-GFP plasmid^55^ by BbsI restriction digest and ligation to generate pSpCas9(NCAPH-3’UTR gDNA)-2A-GFP plasmid. To engineer the template, pMK287 and pMK288 were digested with BamHI to generate mAID-Hygro and mAID-Bsr fragments. Each of these fragments were assembled by Gibson Assembly (NEBuilder^®^ HiFi DNA Assembly, NEB) with 5’ (299bp) and 3’ (293bp) homology arm fragments with 30 bp sequence homology to adjacent assembly fragments on each end and pBS_II+ vector backbone digested with EcoRI. The homology arms were PCR amplified from each side of the Cas9 cut site in the NCAPH 3’UTR (5’ and 3’ Arm forward and reverse primers in ED Tab. 3) using genomic HCT116 DNA as template. These assemblies generated plasmids with mAID-Bsr and mAID-Hygro in between NCAPH 5’ and 3’ homology arms. Template plasmids were transfected into the parental HCT116 TET- OsTIR1 cell line together with pSpCas9(NCAPH-3’UTR gDNA)-2A-GFP using Neon Electroporation System (Thermo-Fischer Scientific). Correct NCAPH-mAID clones were isolated by antibiotic selection using 125 μg/mL hygromycin and 7.5 μg/mL blasticidin and confirmed by PCR, sequencing and degradation confirmed by Immunoblotting. In NCAPH- mAID and NCAPH2-mAID cell lines, endogenous H2B was tagged with EGFP like described previously^53^ and modified for CDK1as chemical genetics for G2-M boundary arrest and synchronous release into mitosis^21,22^. The constructs for CDK1as were gifts from Prof. W. Earnshaw (Addgene #118596 and 118597) and from Prof. Z. Izsvak (Addgene #34879). HCT116 NCAPH-mAID endoH2B-EGFP CDK1as cells were further modified by random integration of α-Tubulin-mCherry for ectopic expression using mCh-alpha-tubulin plasmid which was a gift from Gia Voeltz (Addgene #49149). Clones were selected by treatment with 1.5mg/mL G418 and confirmed by microscopy. Flow cytometry was performed like described previously^56^. Chromosome spreads and immunoblotting was performed as described previously^45,53^. Antibodies used for immunoblotting were NCAPH (1:1000, HPA003008, Sigma-Aldrich), NCAPH2 (1:1000, sc-393333, Santa Cruz), GAPDH (1:6000, G9545, Sigma-Aldrich). The sequences of all DNA oligos used in this study are listed in ED Table 3.

### Cell cycle synchronization and depletion of condensins

To induce expression of TIR1 in the HCT116 TET-TIR1 NCAPH/H2-mAID cell lines, 2ug/mL doxycycline (Sigma-Aldrich) was added 16 hours before depletion. Depletion was initiated with the addition of 500μM indole-3-acetic acid (IAA, sc-215171, Santa Cruz). For depletion of condensins specifically from mitosis (CI-pro or CI-meta), 100μM Auxinole^57^ (MedChemExpress) was added for 16 hours before addition of IAA to prevent potential background depletion from binding of TIR1 to mAID. For synchronization of cell cultures in G2, CDK1as cell lines were arrested in G2 using 0.25 μM 1NMPP1 (Sigma-Aldrich) for 16 hours before release into mitosis, as described previously^23^ and demonstrated in ED Fig. 1g-j. To arrest cells in prometaphase they were treated with 100ng/uL nocodazole (Sigma-Aldrich). For metaphase arrest, proTAME (Bio-Techne) was used at 12.5 μM in combination with 25 μM Apcin (Sigma-Aldrich), like optimized previously^53^. Following arrest in either prometaphase or metaphase, mitotic flush-off was performed like described previously^23^.

### Live cell imaging

HCT116 NCAPH-mAID endoH2B-EGFP CDK1as α-Tubulin-mCherry cells were seeded in poly-L-Lysin coated, 35mm glass bottom μ-dishes (Ibidi) and imaged in a 37°C humidified chamber with 5% CO_2_ on a motorized stage using a ^TM^3i (Marianas Imaging Workstation from Intelligent Imaging and Innovations Inc.) spinning disc confocal microscope equipped with 63X Plan-Apochromat DIC oil objective on inverted Axio Observer Z1 microscope (Zeiss) and CSUX1 spinning disc confocal head (Yokogawa).

### Immunofluorescence microscopy

All immunofluorescence experiments with cells were performed on coverslips coated with poly-L-lysin (P4832, Sigma-Aldrich) following the co-extraction protocol described previously^58^. Antibodies used were NCAPH (1:200, HPA003008, Sigma-Aldrich), NCAPH2 (1:100, sc-393333, Santa Cruz), CENPB (1:200, sc-32285, Santa Cruz), CENPC (1:400, sc-11286, Santa Cruz), α-Tubulin (1:1000, ab18251, Abcam), PICH (1:200, 8886, Cell Signaling). Slides were imaged using a Cell Observer spinning disc confocal microscope (CSU-X1) equipped with a Hamamatsu Orca Fusion camera (C14440-20UP) for Yokogawa spinning disc. These images were then deconvolved using ^TM^Huygens Professional. Images in Fig. 4k and ED Fig. 4k was taken with an LSM 880 Airyscan upright confocal microscope (Zeiss) and images processed using Zen software (Zeiss).

### Native chromosome isolation

HCT116 NCAPH-mAID and NCAPH2-mAID endoH2B-EGFP CDK1as cell lines were transduced with lentivirus to tag TRF1 with BirA and biotinylate the telomere region like described previously^23^. These cells were seeded in 15 cm dishes, arrested in G2 by addition of 1NMPP1 and released for two hours into mitosis with 100 ng/mL nocodazole in the media. Mitotic cells were harvested by “mitotic flush-off” where the medium was flushed 5 times in circular motion over the dish area. This yielded better mitotic purification than traditional “mitotic shake-off” for the HCT116 cell line, which is semi-adherent. Native chromosomes were then isolated from prometaphase cells, as described previously for HCT116 cells^23^. In brief, mitotic cells were hypotonically swollen, lysed in a homogenizer in PA buffer (15 mM Tris-HCl, pH 7.4, 0.5mM EDTA-KCl, 80 mM KCl, 0.1% Tween-20, 1 mM spermidine and 0.4 mM spermine) and the chromosomes purified by centrifugation and glycerol gradient. Unless stated otherwise, all *in vitro* experiments were carried out in PA buffer. All buffers were made with ultrapure water (MilliQ, Millipore) and filtered (0.2 μm pore size; Whatman) before use.

### Optical tweezers experiments

The general workflow for optical tweezers experiments on mitotic chromosomes has been described previously^23,59^. Stretch curves (Fig. 2c,d and ED Fig. 2a) and oscillatory experiments (ED. Fig. 2g-j) were performed on a commercial optical tweezers setup (C-Trap, LUMICKS). Force clamps (Fig. 2h-l) were performed on a similarly equipped, custom-built, setup described in detail previously^23,60^. The distance between the trapped beads was determined by camera tracking. Forces were determined using a position-sensitive detector and binned to the frame rate of distance detection (approximately 20 Hz). Trap stiffnesses were calibrated before each experiment. For stretch curves and oscillatory experiments, they were typically around 0.5 pN/nm. Force clamps were performed at higher laser powers, leading to a larger trap stiffness of around 1 pN/nm. All experiments were performed using streptavidin-coated polystyrene beads (diameter 4.47 or 4.88 µm, Spherotech). For stretch curves, chromosomes were stretched 3 times between a force below 0 pN to a force of 250- 350 pN by moving one of the traps with a constant nominal velocity of 200 nm/s. A linear fit to the trap distance returned an actual velocity of 178±14 nm/s (Mean±Std. Dev.). Since chromosomal stiffness is force-dependent, the chromosome strain rate varied slightly with force. A linear fit to the chromosome length returned an extension velocity of 115±34 nm/s (Mean±Std. Dev.). Only the third stretch was analyzed, as described previously^23^. Oscillatory experiments were performed around a prestress of 50 pN with an amplitude of approximately 100 nm at frequencies of 0.01 Hz, 0.02 Hz, 0.05 Hz, 0.1 Hz, 0.25 Hz, 0.6 Hz, and 1 Hz, as described earlier^3^. For each frequency, at least 3 oscillations were performed, or oscillations were recorded for at least 1 s, whichever was longer. For force clamps, the chromosome was first stretched to a large force between 800 pN and 1000 pN. Then it was clamped alternating at 200 pN and 800 pN, for 4 minutes each, for 3 cycles each.

### Data analysis – optical tweezers

All data analysis was performed using Python with Jupyter notebooks and the Scipy/Numpy framework and MatLab. To load the data generated by the commercial trap, we used the pylake package. The data generated using the homemade trap (.tdms format) were preprocessed in MatLab using the TDMSReader package (Jim Hokanson (2023). TDMS Reader (https://www.mathworks.com/matlabcentral/fileexchange/30023-tdms-reader), saved in a .mat format and then loaded into python using scipy.io.

To calculate the stiffness from chromosome stretch curves, the third stretch curve of a chromosome was selected. Both force and distance were smoothed using a Savitzky-Golay filter with a window size of 81 points and a 3 degree polynomial (scipy.signal.savgol_filter). The stiffness was then calculated by numerical differentiation of the force with respect to the distance. Before calculating mean stiffness-force curves, the individual curves were interpolated to a shared logarithmically spaced force scale. The critical force 𝐹_𝑐_and linear stiffness 𝑘_0_ were extracted as described previously^23^, in brief, curves were interpolated to a logarithmic force scale, and the logarithms of force and stiffness were fitted with a piecewise function 𝑦 = ln(𝑘_0_) for 𝑥 ≤ ln (𝐹_𝑐_) and 𝑦 = 𝑚𝑥 − 𝑚 ln(𝐹_𝑐_) + ln(𝑘_0_) for 𝑥 > ln(𝐹_𝑐_), with the stiffening exponent 𝑚.

To analyze oscillatory experiments, first, the section of data containing oscillations at a given frequency were manually selected. Since force and distance signals were measured and processed by different sensor, it cannot be assumed that the recorded timing is precise. Therefore, in addition to force and distance we also analyzed the bead positions as recorded on the camera, which is in phase with the force acting on the bead and intrinsically synchronized with the distance measure. These three signals (distance, force, and one bead position) were fitted with a sine function *A* sin(2𝜋𝑓𝑡 − 𝜙) + 𝑐 + 𝑚𝑡, with amplitude *A*, frequency 𝑓, time 𝑡, phase 𝜙, a constant offset 𝑐 and linear drift of slope 𝑚. The complex stiffness was then calculated using the amplitudes of the force and distance oscillations *A*_𝐹_and *A*_𝑑_ and the phase of the bead and distance oscillations 𝜙_𝑥_ and 𝜙_𝑑_ as 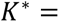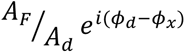

For the analysis of force-clamp data, first the beginning and end points of each force clamp were manually determined as the point when the force stabilized at the target force. To calculate mean curves, first, the measured length was normalized to the length at the beginning of the first force clamp at 800 pN. Then, for each consecutive force clamp at 800 pN, the initial normalized length and time were subtracted from the data (such that the data for each force clamp starts at [0,0]), the data was interpolated to a shared time scale and averages were calculated. To calculate how much irreversible deformation occurred during a force clamp, the differences between the length at the start of the first force clamp at 800 pN and the length at the start of consecutive force clamps at 800 pN were calculated. These values were then normalized by the length at the beginning of the first force clamp at 800 pN.

### AFM experiments

All AFM experiments were performed on poly-L-lysine (PLL)-coated glass coverslips. The coverslips were cleaned in a sonication bath for 1 hour in Hellmanex (2% in ultrapure water), followed by 30 minutes in ultrapure water. After that, the coverslips were dried in an argon flow and stored in a sealed box. Before each experiment, the glass coverslips were activated with air plasma for 3 minutes followed by immediate immersion in ultrapure water for another 5 minutes. The activated slides were then incubated for 30 minutes in a 0.01 % (w/v) poly-L-lysine solution in borate buffer (0.5 mM Na_3_BO_3,_, pH 8.15), at room temperature, followed by thorough rinsing with ultrapure water. Finally, they were dried in a gentle argon flow and placed on the AFM holder for the measurements.

A 10 μl droplet of the chromosome preparation, diluted 1:10 in PA buffer, was spotted on the glass coverslip and allowed to equilibrate for 20 minutes at room temperature before starting the experiments. All AFM experiments were performed on a Bruker Bioscope Catalyst setup coupled with an inverted optical microscope, at room temperature and using the PA buffer as imaging solution. Each chromosome was first located using the fluorescence signal from the inverted optical microscope, then the AFM was used to scan the region of interest and probe the chromosome with higher spatial accuracy and, in the case of AFM force spectroscopy, to run the mechanical characterization. The AFM was equipped with different probes depending on whether it was used for imaging or Force Spectroscopy experiments.

Images were collected in PeakForce Tapping mode, using Bruker SNL-10 probes (C cantilever, nominal tip curvature radius 2-12 nm, nominal elastic constant 0.24 N/m. The force set point and the other imaging parameters were tuned to accurately follow the topological features of the chromosome structure without introducing any deformation.

A Veeco DNP10 probe (cantilever D, nominal tip radius 20 nm and elastic constant 60 pN/nm), calibrated with the thermal noise method^61^ was used for force spectroscopy measurements. Images at lower resolution were used to localize each chromosome; after that, multiple indentations were performed along lines following the shape of the two chromatids. At the end of the indentations, a second image was performed to assess potential deformations or alterations induced by the probing process to the chromosomal structure.

### Data analysis – AFM

Image analysis was performed using Gwyddion 2.58^62^. The recorded force curves were processed with custom made Python scripts to extract the information about the mechanical response of metaphase chromosomes (scripts and a detailed description can be found in the following repository: 10.5281/zenodo.15101776). To consider only indentations performed on the central part of the chromosome body, force curves presenting a contact point < 200 nm from the surface were discarded from the analysis. For extracting the Young Modulus from each force curve, the modified Hertz fit from Dimitriadis et al. ^27^ was applied to the approach curve. Therefore, each curve was fitted from the contact point to an indentation depth equal to 30% of the chromosome thickness (estimated from the distance between contact point and glass surface). The viscoelasticity index 𝜂 is related to the energy dissipated during the indentation process and is defined as 𝜂 = 1 − *A*_𝑟𝑒𝑡𝑟𝑎𝑐𝑡_/*A*_𝑖𝑛𝑑𝑒𝑛𝑡_, where *A*_𝑖𝑛𝑑𝑒𝑛𝑡_and *A*_𝑟𝑒𝑡𝑟𝑎𝑐𝑡_ are the areas under the approach and retraction curves, calculated within the loading and unloading regimes, respectively (i.e., where the tip-sample interactions result in positive force values)^28^. *A*_𝑖𝑛𝑑𝑒𝑛𝑡_ corresponds to the areas shaded in light and dark blue combined in Fig. 1h and *A*_𝑟𝑒𝑡𝑟𝑎𝑐𝑡_ to the area shaded in darker blue.

### Statistics

For statistical difference testing on most of the biophysical data, parametric T-tests for unpaired two-tailed observations were used (function ttest_ind in scipy.stats). The non-parametric Mann-Whitney U Test was used for calculating the p-value of the viscoelasticity index distributions in the AFM results on ΔCI-Pro chromosomes, because the control data did not fit a gaussian distribution. Box-plots were created using the seaborn python package. The box indicates the quartiles, whiskers indicate the range of the whole distribution without outliers. Statistical difference testing on cell biological data was performed using parametric t-tests for two-tailed, gaussian distributed, unpaired data.

### Data availability

All cell biological data generated in this study are provided within the manuscript and its supplemental information. Should any raw data files be needed in another format they are available from the corresponding author upon request. Datasets generated using optical tweezers and AFM are accessible at the public repository Zenodo at this DOI:10.5281/zenodo.15101776 .

### Code availability

All custom code generated and used in this study is available through the public repository Zenodo at this DOI: 10.5281/zenodo.15101776.

## Extended data

**Extended data figure 1.**
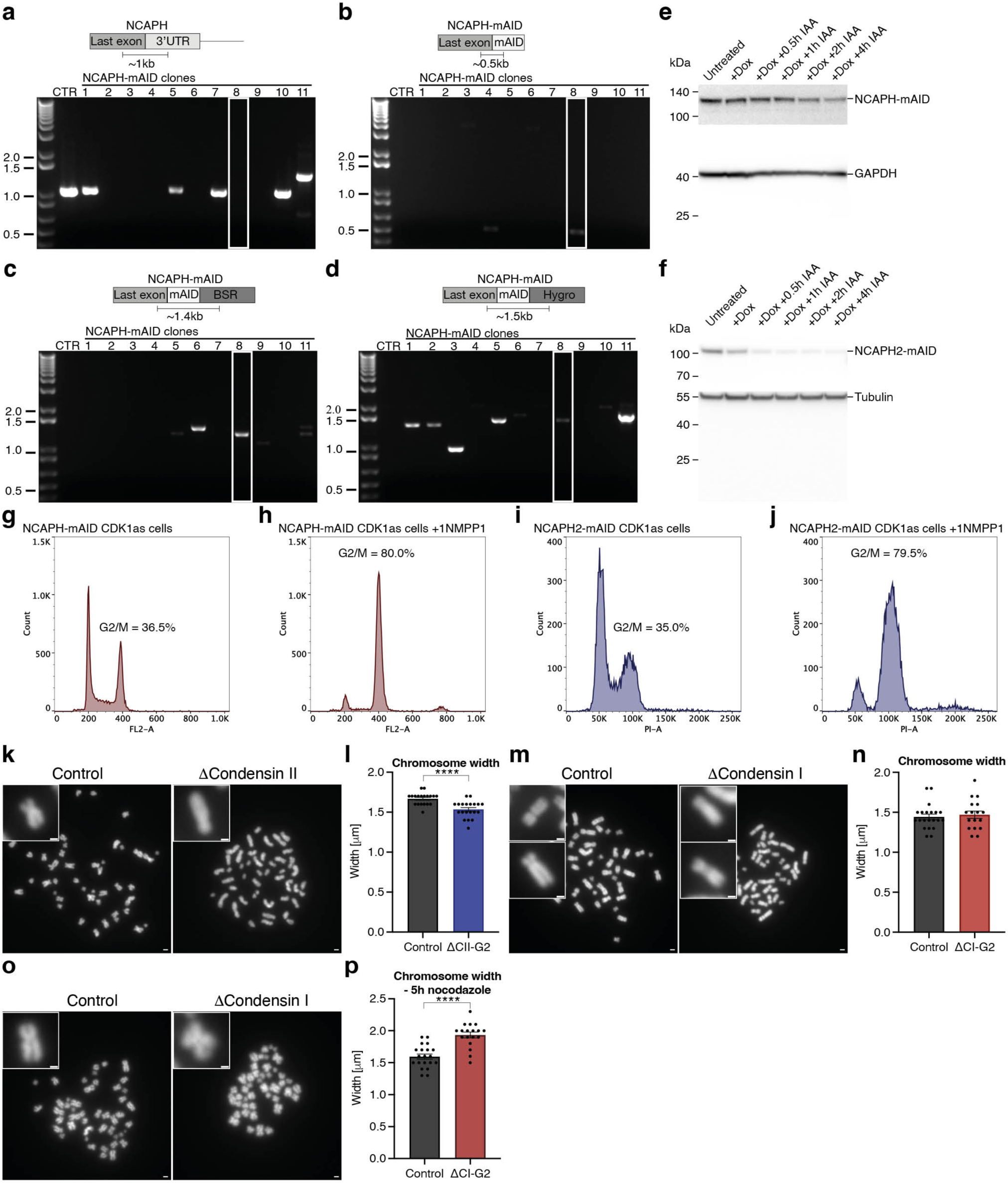
The NCAPH-mAID cell line **a**,**b**,**c**,**d** PCR analysis of genomic DNA from potential NCAPH-mAID clones amplifying the overlaps between the final NCAPH exon and either the 3’UTR (**a**), the mAID tag (**b**), the mAID tag and the blasticidin cassette (**c**), the mAID tag and the Hygromycin cassette (**d**). The white boxes highlight a homozygote NCAPH-mAID clone. **e** Immunoblot of NCAPH depletion from NCAPH-mAID cells following 16 hours of exposure to doxycycline and treatment with 500 μM IAA for 0.5, 1, 2 or 4 hours. **f** Immunoblot of NCAPH2 depletion from NCAPH2-mAID cells following 16 hours exposure to doxycycline and treatment with 500μM IAA for 0.5, 1, 2 or 4 hours. **g**,**h**,**i,j** Cell cycle profiles of NCAPH-mAID CDK1as and NCAPH2-mAID CDK1as cells grown with or without 0.25μm 1NMPP1 for 16 hours**. k**,**l,m,n** Representative images (**k,m)** and quantification (**l,n**) of chromosome spreads from NCAPH2-mAID (**k,l**) or NCAPH (**m,n**) cells treated as in Fig. 1a. **o,p** Representative images (**o)** and quantification (**p**) of chromosome spreads from NCAPH-mAID treated as in Fig. 1a except that they were released into nocodazole induced prometaphase arrest for 5 hours. **k,m,o** boxes in upper left corners show representative chromosomes. **l,n,p** Each datapoint represents the average of a chromosome spread from one cell. Similar results were obtained from two independent experiments. Error bars are SEM. **** p < 0.0001.

**Extended data figure 2.**
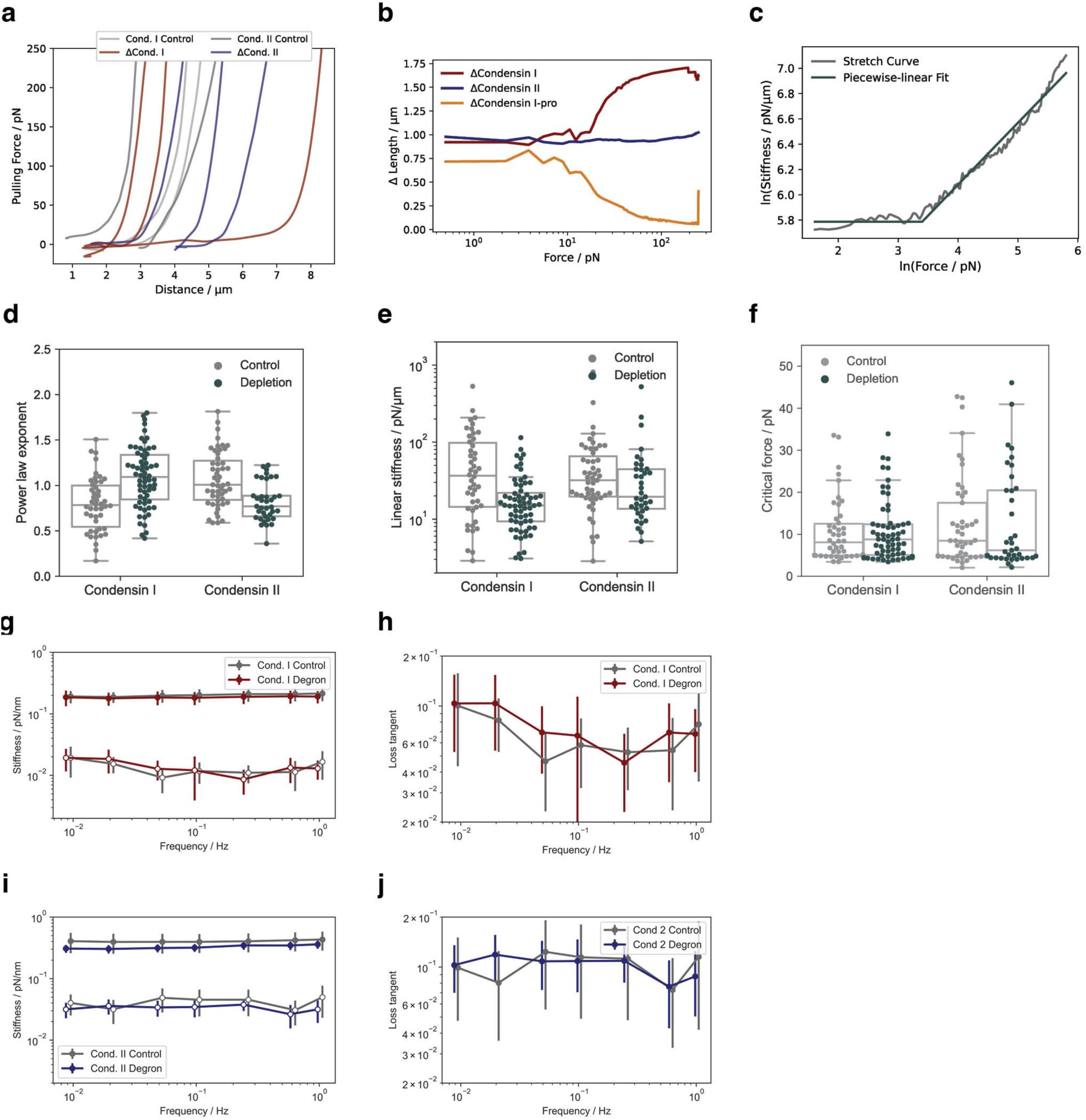
Additional analysis of longitudinal chromosome stretch curves. **a** Selection of example force distance curves of chromosomes depleted of Condensin I or II and the respective controls. **b** The difference between the mean length of degron and control chromosomes at a given force. Most notably, early depletion of Condensin I led to a strong, force dependent length increase. **c** Example fit of a stepwise linear function to the stiffness as a function of the force to extract linear stiffness (**d**), power law exponent (**e**) and critical force (**f**)**. g-h** Frequency dependent complex stiffness for chromosomes depleted of Condensin I (**g**) or II (**h**). **i-j** Frequency dependent loss tangent (imaginary stiffness/real stiffness) for chromosomes depleted of Condensin I (**i**) or II (**j**). Error bars represent SEM.

**Extended data Fig. 3.**
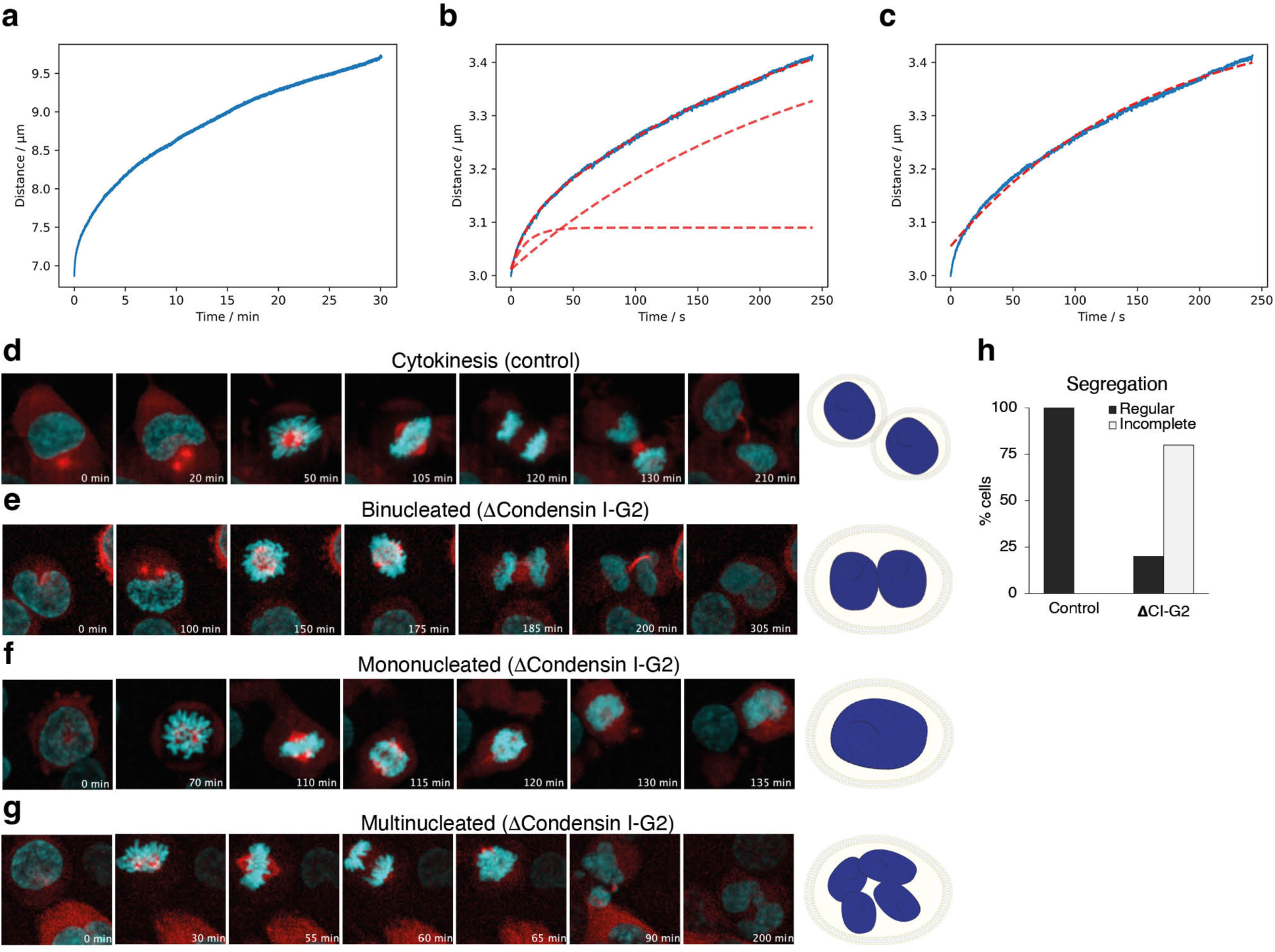
Additional force clamp data and live cell images **(a)** Force clamps over 30 min at 800 pN, even after 30 min no saturation of the length of the chromosome was reached **(b)** Individual force clamp at 800 pN fitted with a double exponential function. The contribution of the two exponentials is also shown individually demonstrating that the slow exponential contributes mostly to the chromosomes elongation **(c)** Same trace as in (**b**) fitted with a single exponential, showing a poor fit. This indicates that chromosomes do not follow a single relaxation time. The two relaxation times were on the order of ∼10 s and ∼150-240 s for all chromosomes, with the longer timescale dominating the total elongation This analysis also revealed that Condensin I depletion in G2 increased the amplitude during both relaxation time periods, without impacting on the timescale itself. **d-g** Representative live cell image series of control (d) and ΔCondensin I cells becoming binucleated (**e**), mononucleated (**f**) or multinucleated (**g**). **h** Quantification of the number of control and ΔCondensin I cells that segregate regularly or incompletely in live cell imaging experiments (control n=23, ΔCondensin I n=30).

**Extended data figure 4.**
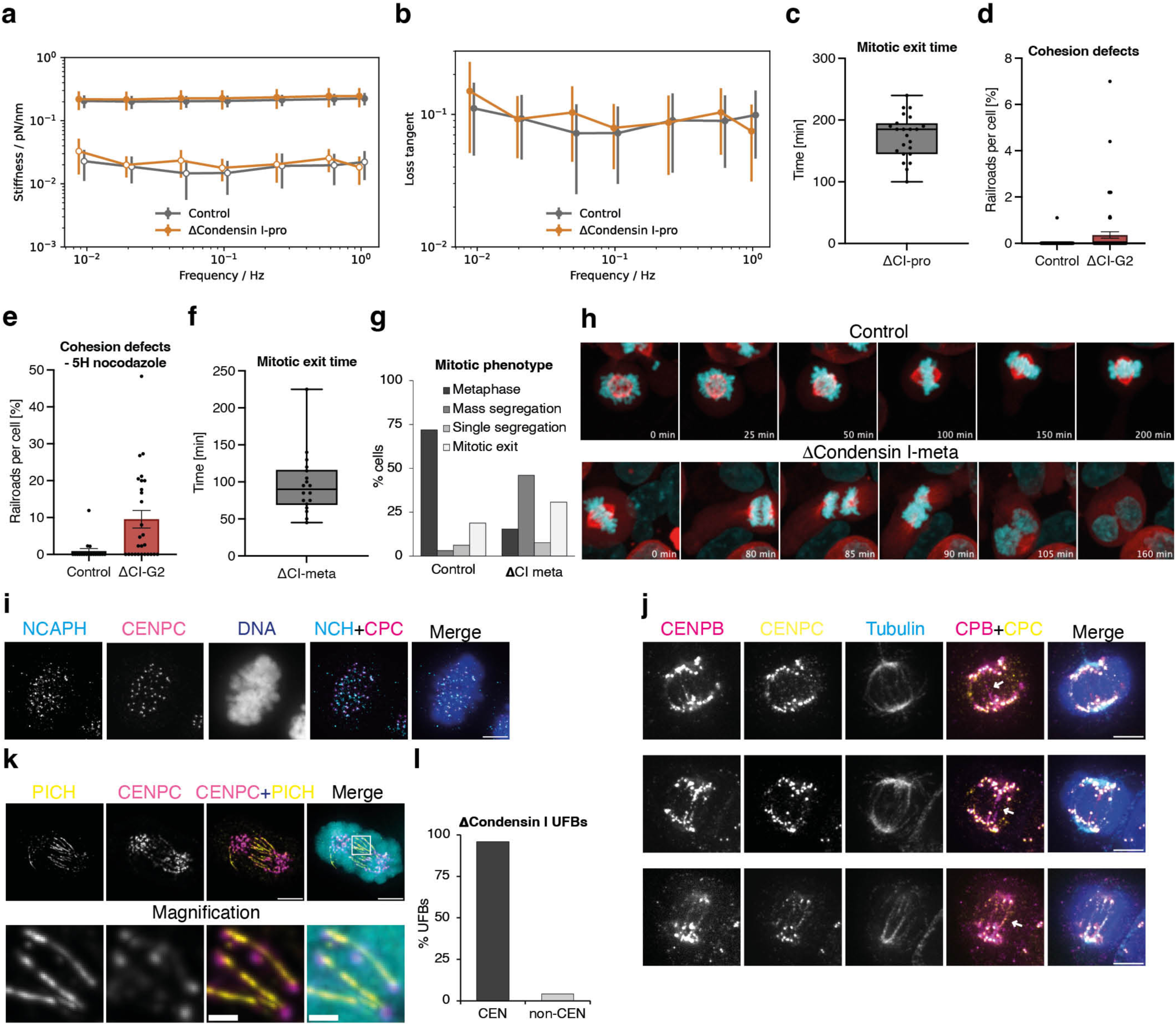
Depletion of Condensin I from cells in mitosis. **a,b** Frequency dependent complex stiffness (**a**) and loss tangent (imaginary stiffness/real stiffness) (**b**) for control and ΔCondensin I-pro chromosomes. **c** Time of mitotic exit from cells in nocodazole prometaphase arrest depleted of Condensin I. Addition of auxin was at time 0. **d,e** Quantification of Railroad chromosomes per cell in chromosome spreads from ΔCondensin I-G2 cells released into nocodazole for 2h (**d**) or 5h (**e**). Each datapoint represents the average of a chromosome spread from one cell. Similar results were obtained from two independent experiments. **f** Time of mitotic exit of cells in proTAME-Apcin induced metaphase arrest depleted of Condensin I. Addition of auxin was at time 0. **g** Quantification of mitotic phenotype of cells leaving metaphase arrest before abandoning cytokinesis. **h** Representative live cell images of control and ΔCondensin I-meta cells arrested in proTAME-Apcin. **i**,**j** Representative images from immunofluorescent staining of NCAPH (**i,** magenta), CENPC (yellow), CENPB (**j**, magenta), Tubulin (**j**, cyan) and DNA stained with DNA (blue) in control cells (**i**) or cells depleted of Condensin I for 2h (**j**) from proTAME-Apcin arrest. The white arrows in **j** highlight stretched centromeric chromatin between sister centromeres. White scale bars are 5 μm. **k,l** Representative images (**k**) and quantification (**l**) of centromeric origin of UFBs in anaphase in NCAPH-mAID cells arrested in prometaphase with Nocodazole for 2h, arrested for 2 more hours with and without Condensin I depletion and then released from prometaphase arrest for 45 min with or without Condensin I depletion. DNA (cyan) is stained with DAPI.

**Extended data Table 1.** Descriptive statistics of the mechanical characterization of chromosomes depleted of Condensin I or II. Rows in black relate to OT results, while rows in green to AFM measurements. For the Young’s Modulus and Viscoelastic Index, we report the statistics related to all the force curves collected in the AFM-FS experiments. Due to the large amount of data, average ± SEM of YM/eta were taken for each chromosome in order to facilitate the analysis and comparison of the obtained results (as presented in the main text). The number of observations for the Youngs modulus and Viscoelastic Index are reported as the number of indentation curves/number of chromosomes.

|  | Degron |  |  | Control |  |  | p-value |
| --- | --- | --- | --- | --- | --- | --- | --- |
|  | Mean | Std Dev | n | Mean | Std dev | n |  |
| **Condensin I-G2** |  |  |  |  |  |  |  |
| Linear Stiffness / pN/μm | 21.3 | 20.3 | 64 | 70.7 | 92.4 | 46 | 7.7E-05 |
| Critical Force / pN | 10.6 | 7.1 | 64 | 10.5 | 7.5 | 46 | 0.96 |
| Stiffening Exponent | 1.1 | 0.3 | 64 | 0.8 | 0.3 | 46 | 4.6E-06 |
| Youngs modulus / kPa | 20.2 | 11.87 | 2348/22 | 15.71 | 10.69 | 2593/20 | 0.087 |
| Viscoelastic Index | 0.32 | 0.09 | 2386/23 | 0.2 | 0.1 | 2582/20 | < 2.2E-308 |
| **Condensin II-G2** |  |  |  |  |  |  |  |
| Linear Stiffness / pN/μm | 49.0 | 89.2 | 37 | 63.6 | 118.2 | 51 | 0.6 |
| Critical Force / pN | 8.4 | 7.2 | 37 | 10.5 | 9.8 | 51 | 0.27 |
| Stiffening Exponent | 0.9 | 0.3 | 37 | 0.9 | 0.3 | 51 | 0.99 |
| Youngs modulus / kPa | 10.55 | 4.39 | 2524/21 | 11.49 | 4.56 | 3308/21 | 0.16 |
| Viscoelastic Index | 0.32 | 0.10 | 2557/22 | 0.31 | 0.09 | 3361/21 | 0.36 |
| **Condensin I-pro** |  |  |  |  |  |  |  |
| Linear Stiffness / pN/μm | 51.3 | 67 | 57 | 44.1 | 59.8 | 44 | 0.58 |
| Critical Force / pN | 11.5 | 10.4 | 57 | 16.5 | 16.0 | 44 | 0.06 |
| Stiffening Exponent | 0.9 | 0.3 | 57 | 1.08 | 0.3 | 44 | 0.0003 |
| Youngs modulus / kPa | 187.88 | 166.84 | 550/12 | 182.14 | 162.47 | 588/11 | 0.56 |
| Viscoelastic Index | 0.40 | 0.16 | 756/12 | 0.50 | 0.15 | 676/11 | 1.24E-32 |

**Extended data Table 2.** Parameters *c*, *A*_1_, *A*_2_,*t*_1_, and *t*_2_ of the best fit of 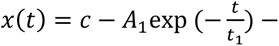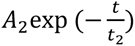 to the mean normalized chromosome length *x* as a function of time *t* at a constant force of 800 pN.

| | $c$ | $A_1$ | $A_2$ | $t_1 / \text{s}$ | $t_2 / \text{s}$ |
| --- | --- | --- | --- | --- | --- |
| **Condensin I-G2 Control** | 0.099 | 0.087 | 0.011 | 240 | 6.6 |
| **ΔCondensin I-G2** | 0.16 | 0.13 | 0.027 | 206.4 | 7.2 |
| **Condensin II-G2 Control** | 0.10 | 0.085 | 0.016 | 239.1 | 9.98 |
| **ΔCondensin II-G2** | 0.11 | 0.091 | 0.014 | 239.8 | 9.9 |
| **Condensin I-pro Control** | 0.13 | 0.099 | 0.029 | 185.2 | 7.9 |
| **ΔCondensin I-pro** | 0.099 | 0.084 | 0.013 | 212.3 | 9.8 |

**Extended data Table 3.**
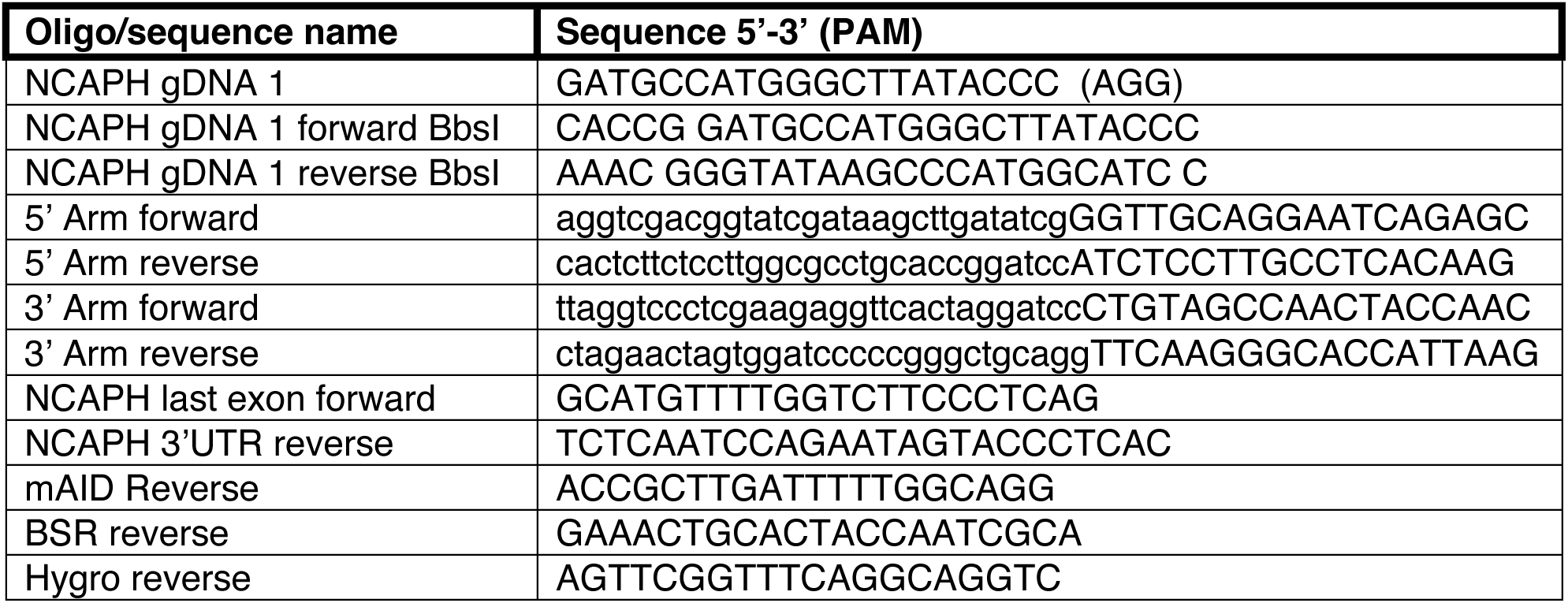
List of oligos used in the study. Small letters denote the non-binding region of primers.

## Supplemental movie captions

**Supplemental movie 1** HCT116 CDK1as NCAPH-mAID H2B-EGFP αTubulin mCherry cells were arrested in G2 for 16 hours then released into mitosis and imaged by timelapse live cell imaging. The cell successfully completes mitosis.

**Supplemental movie 2** HCT116 CDK1as NCAPH-mAID H2B-EGFP αTubulin mCherry cells were arrested in G2 for 16 hours, with depletion of NCAPH in the final four hours then released into mitosis and imaged by timelapse live cell imaging with continued depletion of NCAPH. The cell becomes binucleated and tetraploid following failed mitosis.

**Supplemental movie 3** HCT116 CDK1as NCAPH-mAID H2B-EGFP αTubulin mCherry cells were arrested in G2 for 16 hours, with depletion of NCAPH in the final four hours then released into mitosis and imaged by timelapse live cell imaging with continued depletion of NCAPH. The cell becomes mononucleated and tetraploid following failed mitosis.

**Supplemental movie 4** HCT116 CDK1as NCAPH-mAID H2B-EGFP αTubulin mCherry cells were arrested in G2 for 16 hours, with depletion of NCAPH in the final four hours then released into mitosis and imaged by timelapse live cell imaging with continued depletion of NCAPH. The cell becomes multinucleated and tetraploid following failed mitosis.

**Supplemental movie 5** HCT116 CDK1as NCAPH-mAID H2B-EGFP αTubulin mCherry cells were arrested in G2 for 16 hours, then released into mitosis with nocodazole for two hours and imaged by timelapse live cell imaging. The cell stays in the prometaphase arrest.

**Supplemental movie 6** HCT116 CDK1as NCAPH-mAID H2B-EGFP αTubulin mCherry cells were arrested in G2 for 16 hours, then released into mitosis with nocodazole for two hours before the initiation of NCAPH depletion and imaging by timelapse live cell imaging. The cell eventually abandons prometaphase arrest and mitosis, becoming mononucleated and tetraploid.

**Supplemental movie 7** HCT116 CDK1as NCAPH-mAID H2B-EGFP αTubulin mCherry cells were arrested in G2 for 16 hours, then released into mitosis with proTAME-Apcin for two hours and imaged by timelapse live cell imaging. The cell stays in the metaphase arrest.

**Supplemental movie 8** HCT116 CDK1as NCAPH-mAID H2B-EGFP αTubulin mCherry cells were arrested in G2 for 16 hours, then released into mitosis with proTAME-Apcin for two hours before the initiation of NCAPH depletion and imaging by timelapse live cell imaging. The cell eventually abandons metaphase arrest and mitosis, becoming binucleated and tetraploid.

